# Savings in human force field learning supported by feedback adaptation

**DOI:** 10.1101/2021.03.01.433364

**Authors:** James Mathew, Philippe Lefèvre, Frederic Crevecoeur

**Affiliations:** Institute of Communication Technology, Electronics and Applied Mathematics, Université catholique de Louvain, Avenue Georges Lemaitre 4-6 bte, 1348 Louvain-la-neuve, Belgium; Institute of Neuroscience, Université catholique de Louvain, 53 Avenue E Mounier, 1200 Brussels, Belgium

**Keywords:** Sensorimotor control, force field adaptation, adaptive feedback control, readaptation, savings

## Abstract

Savings have been described as the ability of healthy humans to relearn a previously acquired motor skill faster than the first time, which in the context of motor adaptation suggests that the learning rate in the brain could be adjusted when a perturbation is recognized. Alternatively, it has been argued that apparent savings were the consequence of a distinct process that instead of reflecting a change in the learning rate, revealed an explicit re-aiming strategy. Based on recent evidence that feedback adaptation may be central to both planning and control, we hypothesized that this component could genuinely accelerate relearning in human adaptation to force fields during reaching. Consistent with our hypothesis, we observed that upon re-exposure to a previously learned force field, the very first movement performed by healthy volunteers in the relearning context was better adapted to the external disturbance, and this occurred without any anticipation or cognitive strategy because the relearning session was started unexpectedly. We conclude that feedback adaptation is a medium by which the nervous system can genuinely accelerate learning across movements.

**Significance Statement:** Savings describe the ability of healthy humans to faster relearn a previously acquired motor skill. It is debated whether savings result from implicit changes in the learning rate or an explicit strategy. Given recent evidence that feedback control plays a key role in movement adaptation, we hypothesized that this component also contributed to savings in the context of force field adaptation. We confirm that relearning was faster and demonstrate that the very first relearning movement was better adapted to the external disturbance, i.e., the online corrections in the first re-exposure trial carried imprints of feedback adaptation from the previous sessions in the absence of any explicit strategy. We conclude that this non-explicit feedback component can genuinely accelerate motor learning.

## Introduction

Humans adapt to force field perturbations in reaching movements within a few trials. During adaptation, a motor memory of the newly acquired mapping is created, which is fragile in the early stage of adaptation and progressively becomes stable (Krakauer and Shadmehr, 2006). When exposed to the same perturbation a second time, it was documented that relearning is faster and more complete than during the first exposure, a phenomenon called “savings”(Brashers-Krug et al., 1996; Shadmehr and Brashers-Krug, 1997; Caithness et al., 2004; Overduin et al., 2006; Focke et al., 2013; Stockinger et al., 2014; Coltman et al., 2019). There are two leading hypotheses to explain savings: the first is that the predictive, feedforward component of motor adaptation changes faster across trials due to the repeated exposure to similar perturbation (Gonzalez Castro et al., 2014); alternatively, it was also reported that savings reflect an explicit, cognitive strategy expressed following the prior learning experience (Taylor et al., 2014; McDougle et al., 2015). Currently, the question of whether savings reflect changes in learning rate or an explicit strategy remains largely open and is central to understand neural mechanisms underlying human motor adaptation.

A key question in this debate is to understand the role of feedback control. First, it is clear that the time-varying force fields recruit and adapt the feedback control system (Wagner and Smith, 2008; Yousif and Diedrichsen, 2012; Cluff and Scott, 2013; Joiner et al., 2017; Maeda et al., 2018). Additionally, recent studies highlighted the importance of online adaptive control in reaching movements and argued that rapid feedback adaptation contributed to the trial-by-trial adaptation (Crevecoeur et al., 2020b, 2020a; Mathew et al., 2020). Moreover, changes in long-latency feedback responses to perturbations paralleled the fast learning timescale (Smith et al., 2006; Coltman et al., 2019; Coltman and Gribble, 2020), which must be involved if an increase in learning rate occurs over the course of few tens of trials. Altogether the link between feedback control and the fast timescale of motor adaptation warranted an investigation of the potential influence of feedback adaptation in savings during force field learning.

More precisely, we hypothesized that a genuine acceleration during relearning could originate from feedback adaptation, unveiling a novel component of savings to consider. The testable prediction is that if a feedback adaptation component from the first learning session facilitates relearning, it should be evident in the online corrections made in the very first readaptation trial, in the absence of any anticipation or explicit strategy. Note that the current explanation of savings (changes in the feedforward learning rate, or explicit strategies), do not predict any effect within the very-first relearning trial when the perturbation is introduced unexpectedly. Here we demonstrate the existence of a feedback component in savings by designing a force field adaptation task (with unexpected readaptation), that rules out any contribution of an explicit strategy and allows showing an effect of faster relearning even for the first trial of readaptation, for which there is no role of the feedforward component. Our results confirm that feedback control in the first relearning trial is better tuned to the force field, and suggest that this novel feedback-mediated component contributes to savings in human reaching adaptation.

## Materials and Methods

### Experimental design

Sixteen self-declared right-handed healthy adults (age = 24.4±3.5, 4 male) were recruited. The experimental paradigm was approved by the Ethics Committee of the host institution (UCLouvain, Belgium) and complied with the Declaration of Helsinki. Participants were compensated for their participation. Before the experiment, all the participants provided written informed consent.

Participants were requested to sit comfortably in front of a robotic device (KINARM, Kingston, ON, Canada), with their forehead resting on a resting pad. They were instructed to perform visually guided forward reaching movements in the horizontal plane using the handle of a robotic arm, towards a virtual target on the screen in front of them.

There were two types of movement conditions. The first type corresponds to Baseline Trials: These were reaching movements from starting position (a filled circle with a radius of 0.6cm) to a goal position fixed at 15cm from starting position. The direct vision of the arm and hand was blocked, but the cursor aligned to the handle was always visible providing feedback about the current hand position. Participants had to wait at the starting position for a random delay uniformly distributed between 2s and 4s. The goal position was initially presented as an open red circle (radius 1.2cm) and later as a filled red circle, indicating the cue to initiate the movement. For a successful trial, the goal circle became filled with green when the participants reached the target within 600ms to 800ms (including reaction time) and stabilized there for at least 1s. If they moved too fast, the goal circle changed back to an open circle. If the movement was too slow, it remained red. The feedback about timing was provided to encourage consistent movement times, and despite the failure to timing criterion, all movements were included in the analyses. The second type was Force field trials (FF). In these trials, participants experienced a lateral force perturbation proportional to the forward hand velocity (Fx= ±Lẏ, L = 13 Nsm^−1^). Thus, force fields were either clockwise (CW) or counter-clockwise (CCW). Participants were divided into two groups: group G1 experienced only CCW force field during adaptation and readaptation blocks and group G2 experienced only CW force field (Fig 1c).

**Figure 1.**
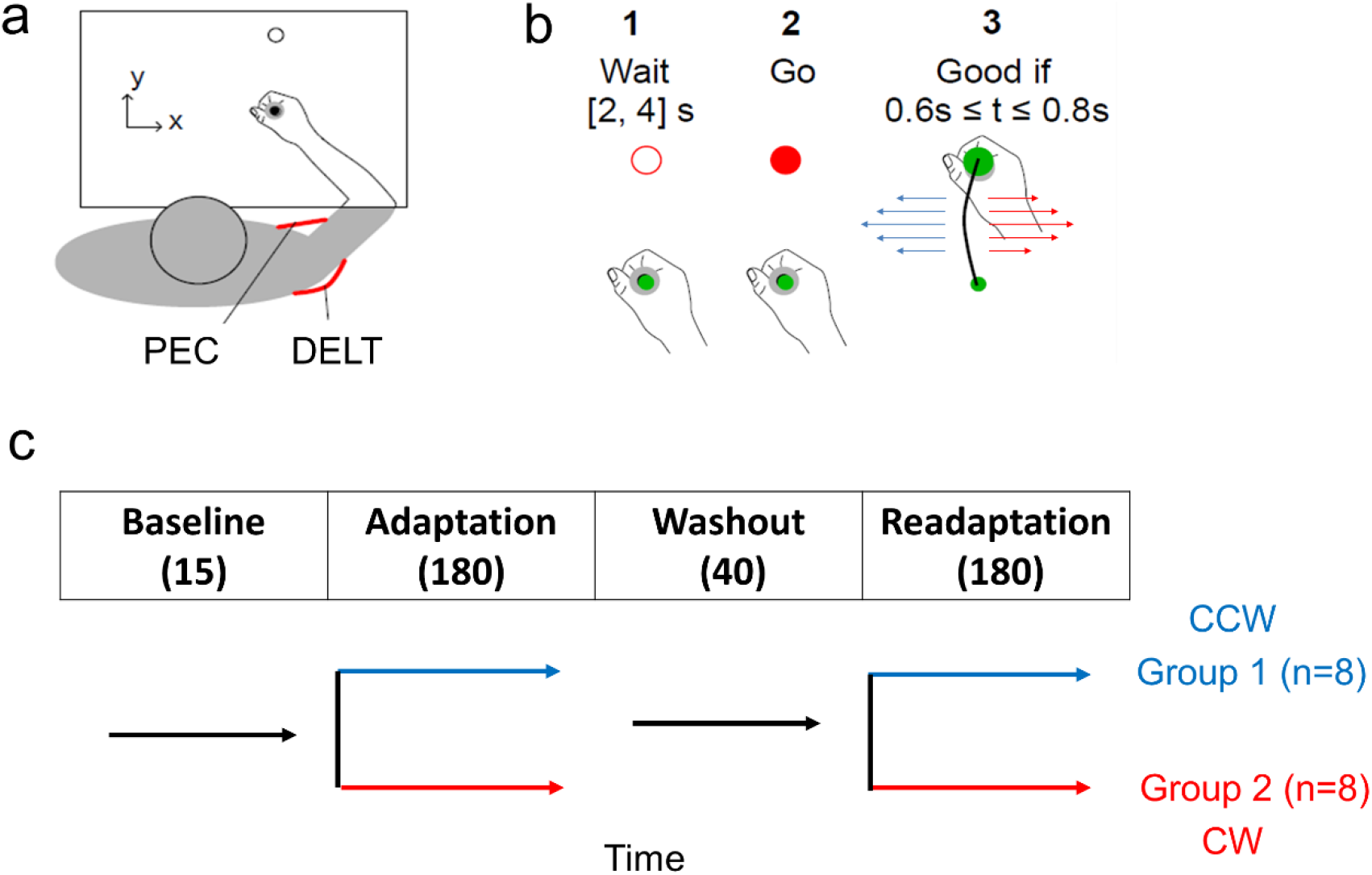
a) Experimental setup, an illustration of the workspace. Electromyogram was recorded from Pectoralis Major (PEC) and Posterior Deltoid (DELT). b) Illustration of a successful reaching trial. Participants were instructed to wait for the go signal and then to reach the goal within 0.6 to 0.8 s of the go signal. Visual feedback about movement timing was provided to encourage participants to adjust their movement speed, but all trials were included in the analyses. Reaching trials were with or without force field perturbation. During perturbation trials, a velocity-dependent force field was applied in either a clockwise (CW) or counterclockwise direction (CCW) (Blue arrows represent CCW and red CW). c) Experimental design. Participants completed 415 trials in a session. Force field was turned off during baseline and washout blocks and turned on during adaptation and readaptation blocks. The first group of participants experienced only CCW (blue timeline, n=8) and the second group only CW force fields (red timeline, n=8). Perturbations were introduced without prior warning.

Experimental sessions consisted of four blocks of trials. The first block was the baseline block with 15 baseline trials; the second block was the adaptation block with 180 FF trials; the third block was the washout block with 40 baseline trials, and the fourth block was the readaptation block with 180 FF trials in the same direction as during the adaptation block. Adaptation and readaptation phases were separated by 40 trials performed in the null environment to washout learning.

### Data analysis and statistics

Signals from the robotic device were sampled at 1000Hz. X and Y coordinates of the cursor/robotic handle and the forces at the interface between the participant’s hand and the handle were digitally low-pass filtered with a fourth-order, dual-pass Butterworth filter with a cut-off frequency of 50Hz. Velocity signals were obtained from numerical differentiation of position signals (4th order, finite difference algorithm). Electromyogram (EMG) was recorded using Bagnoli Desktop System (Delsys, Boston, MA, US) at 1000Hz sampling frequency and digitally band-pass filtered (4th order dual-pass: [10, 400] Hz), rectified, then averaged across trials and participants. EMG electrodes were positioned over the muscle belly of the shoulder flexor - Pectoralis Major (PEC) and shoulder extensor - Posterior Deltoid (DELT), the main muscles recruited when performing lateral corrections against the force field perturbations used in this experiment, and on the right foot ankle for ground electrode.

Three events were used as timing references. 1) The Go cue to initiate the movement, 2) reach onset which was defined as the moment when the cursor exited the home target 3) trial end when the cursor stabilized at the goal position. Hand paths were averaged across the subjects. For each subject, the lateral (v_x_) and forward (v_y_) component of the peak velocity was computed for each trial. We followed standard EMG normalization procedures as follows. We performed two calibration blocks, in which participants were requested to counteract a 10N force towards left or right (3 trials in each direction), activating DELT and PEC, respectively. We averaged EMG during the stationary phases of calibration trials (1s time window) to get a reference value for each muscle. The EMG trials during the experiment were normalized to this reference value. The normalized EMG traces during the first trials of the adaptation and readaptation blocks were averaged across subjects and were aligned to the reach onset. For statistical comparison, for each subject, each EMG trace was averaged within a 100ms window before reach onset.

The evolution of hand movement variation across the trials was assessed by means of exponential fits on parameters like path length, maximum lateral displacement, maximum lateral target overshoot, peak terminal force, initial reach angle and the correlation between commanded and measured forces. The path length and maximum lateral deviation are related, and they are minimum for straight movements. The lateral target overshoot reflects the absolute magnitude of the interaction force at the handle near to the target, ie, the peak terminal force and they are associated with adaptation of feedback control (Crevecoeur et al., 2020b, 2020a; Mathew et al., 2020), and the correlation between commanded and measured forces was also previously demonstrated as an indicator of adaptation. The exponential model has the following form:

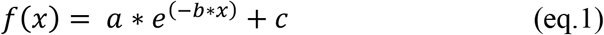

and we regressed the 3 free parameters a, b and c, which correspond to the amplitude of adaptation change, adaptation rate and asymptotic performance, respectively. The exponential models were fitted in the least square sense and the significance of the fit was determined based on whether the 99.5% confidence interval of the exponent responsible for the curvature of the fit included or not the value of zero (p < 0.005). The nonsignificant fits were associated with p >0.05. R^2^ statistics were derived as follows:

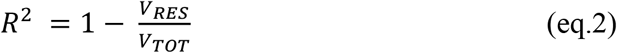

where V_RES_ denotes the variance of the residuals and V_TOT_ is the total variance of the data (James et al., 2013; Crevecoeur et al., 2020b). Repeated measures ANOVA was used to document statistical differences between adaptation and readaptation blocks in CW and CCW perturbations with a significance level of p<0.05. For ANOVAs, effect sizes are reported as partial eta squared 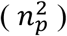(Fritz et al., 2012; Lakens, 2013).

For several analyses as specified in the Results sections, we were interested in confirming the absence of any difference between the two datasets. The suitable approach to provide this information is to report Bayes’ factors, which compares the ratios of the likelihood of observing the two samples and can provide information about whether there is any evidence in favor of either assumption (presence or absence of difference) (Rouder et al., 2009, 2012; Faulkenberry, 2018). Bayes factors were used to characterize non-significant effects, based on the likelihood of two models (means are equal, vs means are different). When we observed non-significant effects, we calculated BF01, standing for the ratio between the likelihood of the data under H0 (there is no effect of the tested predictor), and H1 (there is an effect of the predictor). With this definition, BF01 is the evidence supporting H0, and we use the following interpretation table: BF01 between 1 and 3: only worth mentioning; BF01 between 3.2 and 10: substantial evidence; BF01 >10: strong evidence.

Force field adaptation was assessed using the correlation between the commanded force and the measured force. The commanded force in the environment was calculated offline as the forward velocity multiplied by the scaling factor that defines the FF (L = ±13 Ns/m). The measured force is the force at the interface between the hand and the robotic handle, which is measured with the force transducer. The differences between these two forces were well correlated with the lateral acceleration (For a detailed description see (Crevecoeur et al., 2020b)). The initial reach angle (initial movement direction) was calculated as the angle between reach movement and the straight line connecting starting and target position, a positive value indicates an angle towards the left of this straight line.

## Results

Our analyses will proceed as follows, we first confirm the presence of savings by showing that readaptation was indeed faster than adaptation. Second, we demonstrate that the very first trial in the readaptation block was already better adapted after verifying that trials at the end of the washout phase were indistinguishable from the baseline trials, and without any explicit strategy as the readaptation blocks were introduced unexpectedly. Third, we highlight the feedback control component that contributes to this better readaptation performance and savings. Finally, we present control analyses to verify the absence of anticipation at the beginning of the readaptation phase and to rule out the presence of any non-specific strategy linked to the co-contraction strategy that leads to this better performance.

Reach movements were first strongly impacted by the introduction of the perturbations and then became smoother with practice (Fig 2a). To quantify the change in reach performance within and across blocks, we extracted few key parameters 1) the total distance travelled by the hand cursor (pathlength), 2) the maximum lateral hand deviation along the direction of the force field 3) the lateral target overshoot near the end of the movement 4) the peak terminal measured force and 5) the correlation between commanded and measured forces. In all these parameters, it was visible that the initial trial in the readaptation block was different from the very first trial from the adaptation block, and their values were closer to those observed from adapted movements in the first block.

**Figure 2.**
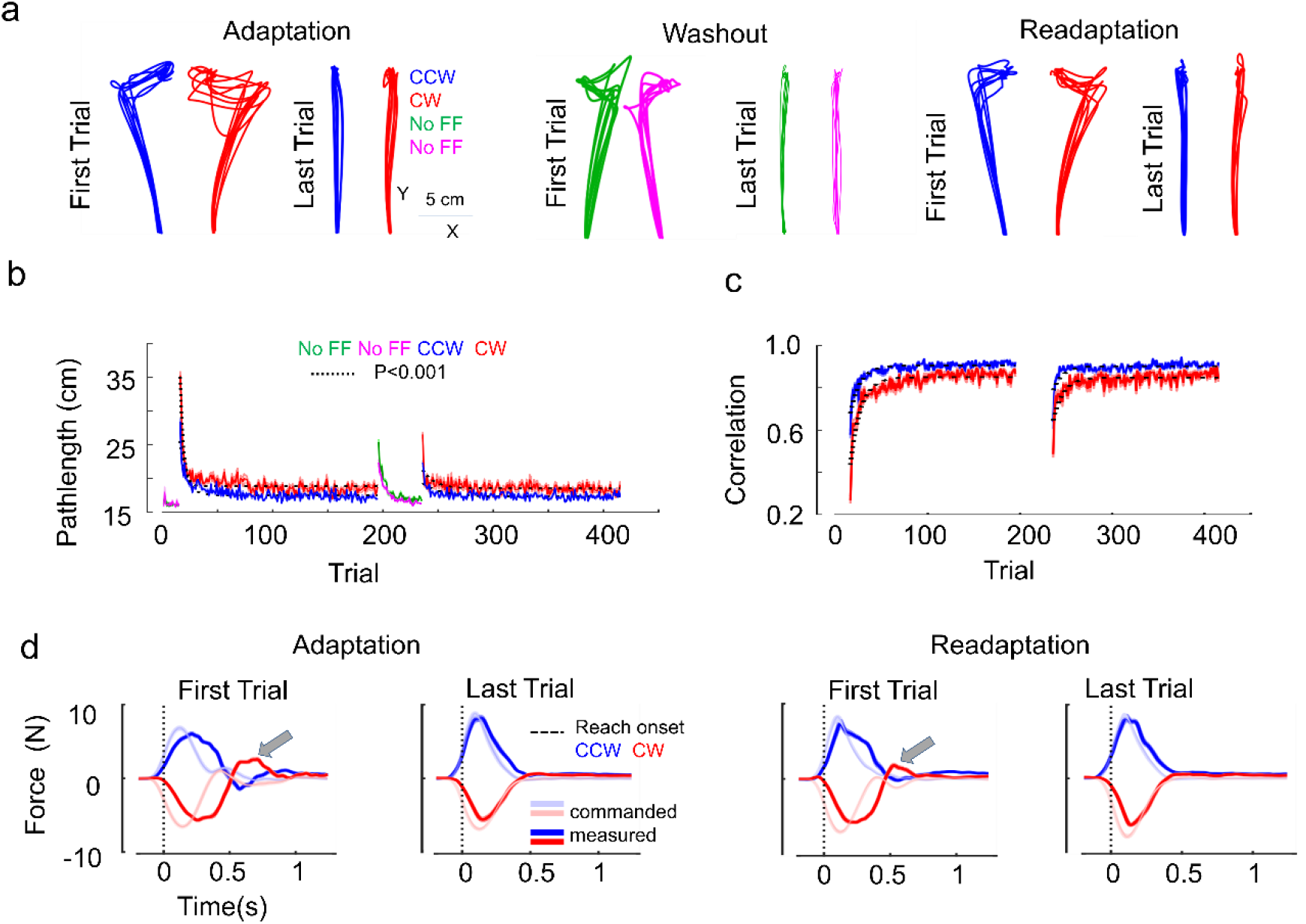
a) Hand paths from all participants for the first and the last trial in adaptation, washout and readaptation blocks. CCW and CW FF trials are depicted in blue and red, respectively. No FF trials during washout block for group 1 and group 2 are depicted in green and magenta respectively. The last trials in adaptation and readaptation blocks are well adapted to the FF, with reduced lateral deviation and target overshoot. The first trial in washout block shows after-effects. The last trial in the washout block is shown to highlight visually that the participants returned to baseline performance by the end of that block. b) Pathlength for No FF, CCW and CW FFs across trials. The solid lines indicate strongly significant exponential fits (p<0.001) that reveal curvature across force field trials. Shaded areas around the mean represent the SEM across participants. Adaptation (trial no:16-195) and readaptation (trial no: 236-415) blocks are contrasted with no force field trials ie, baseline (trial no: 1-15) and washout (trial no:196-235). c) Correlation (R^2^) between commanded perturbation force and the interaction force produced at the interface between participants’ hand and the handle (measured force), across trials. The evolution of changes in force profiles across trials is evaluated using the correlation, better correlation means better compensation for the applied force. d) Commanded (light colour) and measured forces (dark colour) for the first and last trial of adaptation and readaptation blocks. CW in red and CCW in blue. The vertical dotted line represents the reach onset. The second peak force (highlighted with grey arrows) is nearly flattened for the last trials. Importantly, the first trial in the readaptation block shows better force control compared to the first trial in the adaptation block. The increased correlation and the better tuning of force profiles across trials are indications of better online feedback control.

For pathlength, a significant exponential decay across trials was observed in the adaptation block (Fig. 2b, p<0.001, R^2^ = 0.80 for CW and R^2^ = 0.71 for CCW FFs) and in the readaptation block (p<0.001, R^2^ = 0.30 for CW and R^2^ = 0.37 for CCW). We calculated the time as the number of trials to reach 90% of the asymptote of the learning curve and found that relearning was faster than learning by 75% in CCW (29 versus 7 trials to reach 90% of asymptote for Ad and Re respectively) and by 44% in CW (41 versus 23 trials). The time to reach asymptote was directly related to the rate of increase or decrease of an exponential phenomenon. We privileged this variable over individuals’ learning rates that tended to be more variable across participants.

To follow the analysis on pathlength presented above, we evaluated the correlation between commanded and interaction forces and observed a significant exponential increase in correlation across trials in the adaptation block (Fig 2c, p<0.001, R^2^ = 0.84 for CW and R^2^ = 0.80 for CCW), as well as in the readaptation block (p<0.001, R^2^ = 0.61 for CW and R^2^ = 0.49 for CCW). Here also we calculated the time to reach 90% of the asymptote of the learning curve and found that relearning was faster than learning by 80% in CCW (10 versus 2 trials to reach 90% of asymptote for Ad and Re) and 36% in CW (25 versus 16 trials). Thus, this analysis confirms that relearning was indeed faster.

Since the commanded force was a function of hand forward velocity, we could interpret the increased correlation across trials as the tuning of hand force with the hand forward velocity (Fig 2d), which is a marker of adaptation (Crevecoeur et al., 2020a, 2020b; Mathew et al., 2020). Thus, both the reduction in pathlength and the increase in correlation reached their asymptotes faster, which highlighted the presence of savings.

In light of the tendency to reach the asymptote faster, our central question was to characterize the very first trial in learning and relearning contexts. We took the pathlength of the first trial in the two blocks and performed repeated-measures ANOVA over the learning phase (Adaptation and Readaptation, after pooling CW and CCW together). We found that the pathlength was significantly reduced in the first trial of readaptation block compared to that of adaptation block (Fig 3a, G1 and G2 pooled mean: Ad=31.8cm vs Re= 25.2cm, 20% reduction, F(1,7)=27.98, p=0.001, 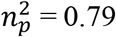). Besides, we compared the correlation between commanded and interaction forces from the same very first trials from each block and found again that the correlation was significantly improved in the very first trial of readaptation block (Fig 3b, G1and G2 pooled mean: Ad=0.42 vs Re=0.60, 43% improvement in Re, F(1,7)=26.24, p=0.001, 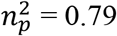).

**Figure 3.**
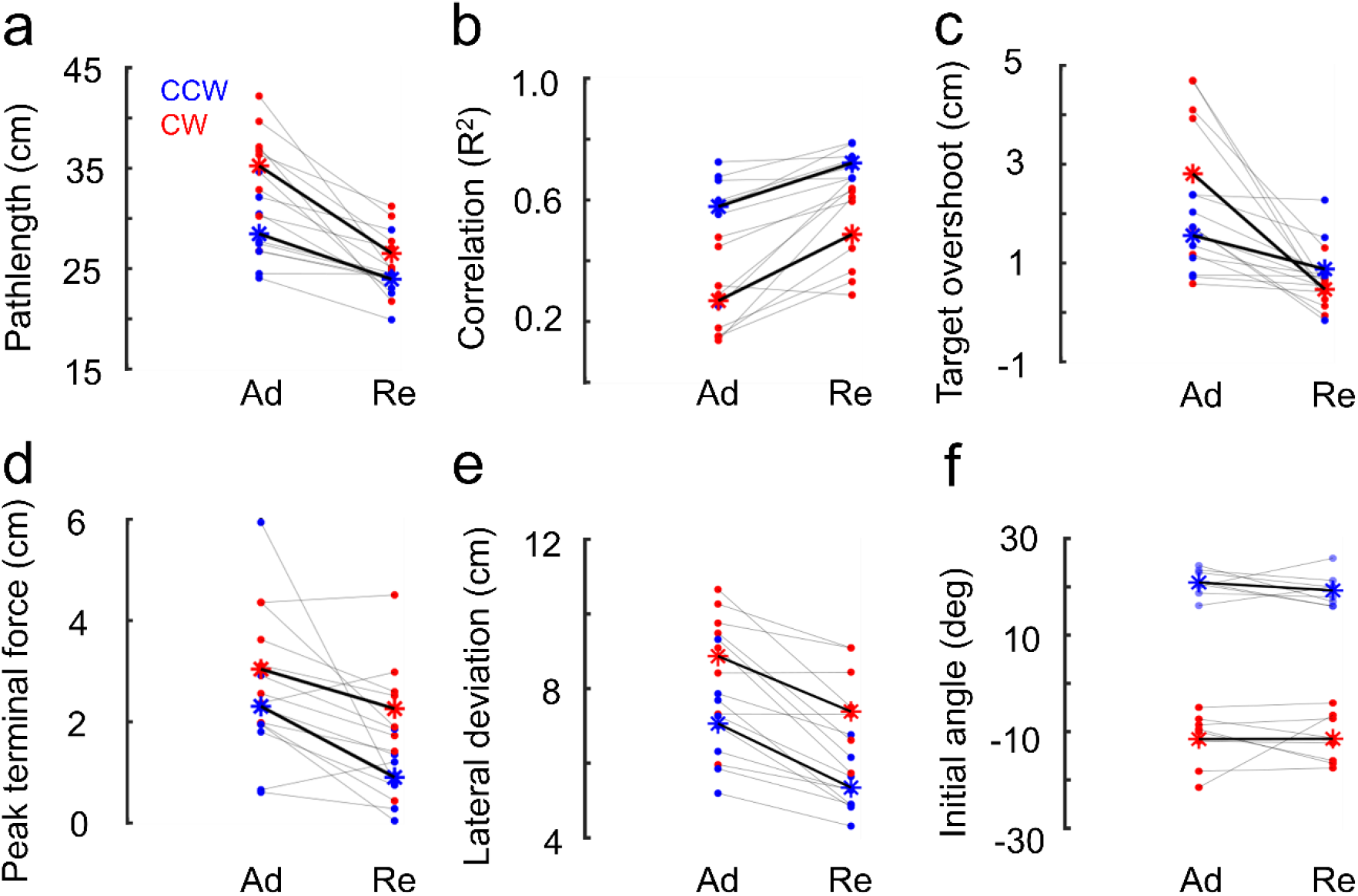
a) Pathlength of the first trial in adaptation (Ad) and readaptation (Re) blocks. Each dot represents one participant, CCW and CW FF trials are depicted in blue and red, respectively. The mean of each group is represented by an asterisk (*). The grey lines represent variations for each participant and the black lines represent the mean variation between Ad and Re for each group. During the first trial in adaptation (Ad) and readaptation (Re) blocks, b) Correlation (R^2^) between commanded and measured force. c) Target overshoot in the lateral direction d) Absolute peak terminal force applied on the handle e) Maximum lateral deviation f) Initial movement direction towards the target (initial angle made by the reaching movement with the straight line connecting starting and target position). Despite the absence of anticipation (comparable initial angle), the reduced path length, target overshoot, peak terminal force, lateral deviation and a higher commanded-measured force correlation in the first readaptation trial highlight the recall of an efficient feedback control response during the very beginning of readaptation phase.

Similar observations (ie, first relearn trial is better) were also made in other parameters characterizing movement kinematics like the maximum lateral hand path deviation (Fig 3e, G1 and G2 pooled mean: Ad=7.9cm vs Re=6.3cm, 20% smaller deviation for the first trial in readaptation block compared to that in adaptation block, F(1,7) = 29.33; p=0.001, 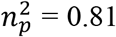), the lateral target overshoot near the end of the movement (Fig 3c, G1 and G2 pooled mean: Ad=2.18cm vs Re=0.67cm, 69% smaller overshoot for the first trial in readaptation block compared to that in adaptation block, F(1,7) = 19.4; p=0.003, 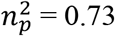), and the absolute peak terminal force (Fig 3d, G1 and G2 pooled mean: Ad=2.67N vs Re=1.58N, 41% smaller peak terminal force during the first readaptation trial, F(1,7) = 12.36; p=0.009, 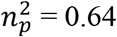). Note that the reduction in terminal force (also in Fig 2d, highlighted in grey arrows) occurred without any change in the force field, since hand forward velocities and therefore lateral forces were similar in the first and last trials of both blocks (rmANOVA, F(7,49)=1.76; p=0.12, 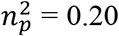; Bayes Factor(BF) =2462, strong evidence for the null hypothesis that the means are equal).

All extracted parameters from the very first relearning trials presented the features of more adapted movements. If we can demonstrate that there was no anticipation of the force field at the beginning of the readaptation phase, that is the washout was complete, then the improvement in the first relearning trials must correspond to an adaptation of online feedback control (Crevecoeur et al., 2020b, 2020a; Mathew et al., 2020). The foregoing analysis shows that it was indeed the case.

To further ascertain the contribution of feedback adaptation in faster relearning, it is thus necessary to verify that movements performed in the null field were similar across the baseline block and the washout block, during which the participant’s reaching movements recovered from the after-effects and returned to baseline performance. The washout block was intended to eliminate the feedforward component about the perturbation and at the end of 40 washout trials, we observed zero anticipation. We compared the last baseline trial with the last washout trial and found no significant difference in both groups in terms of pathlength (F(1,7) = 1.02, p = 0.34, 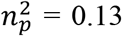; BF =1.64, evidence for the null hypothesis: washout trials and baseline trials were equal) and initial angle (initial movement direction) (F(1,7) = 1.76, p=0.23, 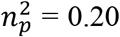; BF =1.15, evidence for the null hypothesis: washout trials and baseline trials were equal). Thus the baseline and late washout trials were statistically indistinguishable, suggesting that deadaptation was complete.

Another way this could be verified was by looking at the initial reach angle from the first readaptation trial: indeed if deadaptation was not complete, we would expect a difference in the initial angle, used as a proxy of feedforward strategies. Again, by comparing the first trial in adaptation and readaptation blocks, we found no significant difference in the initial angle (Fig 3f, G1 mean= 20.9 deg and 19.2 deg, G2 mean = −11.47 deg and −11.5 deg for Ad and Re blocks respectively, F(1,7) =0.47, p=0.51, 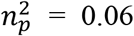; BF = 2.18, evidence for the null hypothesis: aiming angles in both blocks were equal). This confirms that the first trials in both blocks had similar (zero) anticipation, allowing us to rule out a partial anticipatory compensation, or an explicit strategy.

The previous observations confirmed that there was no residual or partial anticipation for the force field, but it remains possible that feedback control was different. To address this possibility, we analyzed the feedback control strategy by looking at the positional variance across trials over the reach time. Because perturbations may elicit changes in the control strategy independent of adaptation (Crevecoeur et al., 2019), we avoided applying disturbances to directly probe feedback gains. Instead, we observed the variance across trials as an indicator of how the natural variability of movements was regulated as a proxy of the feedback controller (Todorov and Jordan, 2002). The reduction in peak terminal force already argued against a default increase in feedback gains, since this would predict an increase in interaction forces. Besides, we computed the positional variance across late washout trials (15 trials) during the reach time (800ms) and compared it with that of baseline trials (15 trials). We did not find any evidence that the positional variance in both x- and y-direction was lower at the end of washout in comparison with baseline trials.

Finally, we investigated whether participants used or not co-contraction to modulate the limb intrinsic properties, and increased their feedback gains (Burdet et al., 2001; Franklin et al., 2008; Crevecoeur et al., 2019), potentially resulting in higher movement accuracy (Gribble et al., 2003). To evaluate the possible contribution of co-contraction to the improved control during the very first readaptation trial, we quantified EMG levels of a pair of agonist-antagonist shoulder muscles known to be strongly recruited by the perturbations that were used in this study (Fig 4 a, b) (Franklin et al., 2008; Crevecoeur et al., 2019). For each participant, we computed the mean EMG activity during a 100ms window before the reach onset for both PEC and DELT muscles (Fig 4c). Then we compared the EMG in baseline trials with the first trial in adaptation and readaptation blocks for each participant and found no significant difference (repeated measures ANOVA for PEC: F(2,30)=0.26;p=0.76, 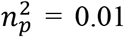;BF =24.31 and for DELT: F(2,30) =0.61, p=0.54, 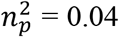; BF = 16.91, strong evidence for the null hypothesis that muscles activities are equal). This supports the argument that the better reach control in the first trial in readaptation blocks was not due to any co-contraction modulating the limb intrinsic properties, instead, it involved a feedback adaptation component that was described previously (Crevecoeur et al., 2020a; Mathew et al., 2020).

**Figure 4.**
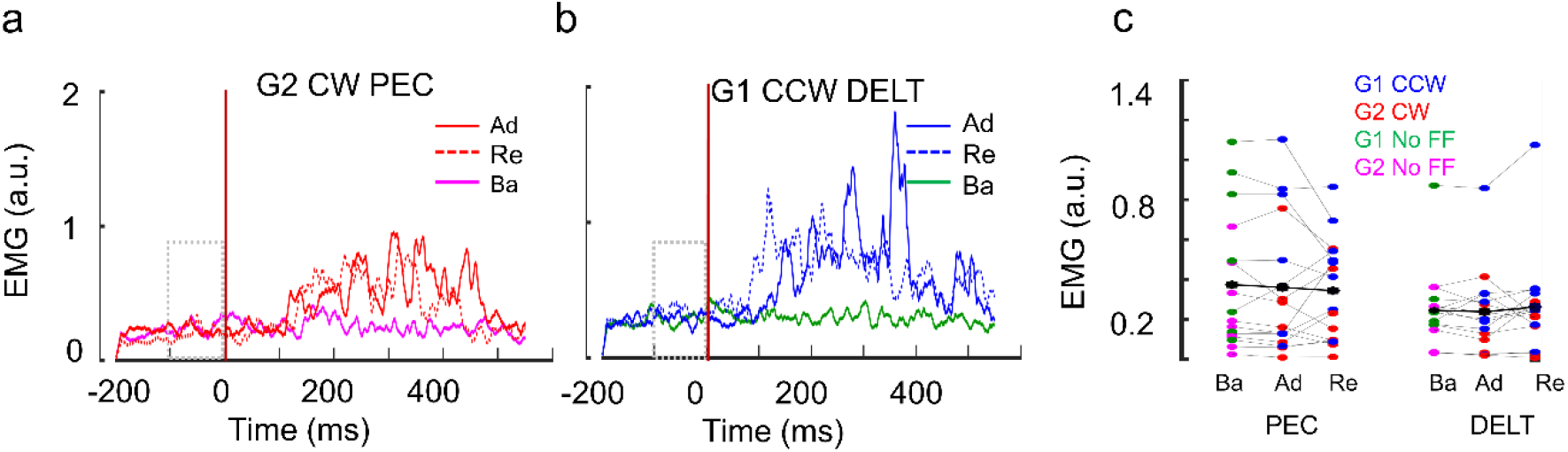
a) Normalized EMG activity in pectoralis major muscle during CW perturbation (group 2). Each red EMG trace corresponds to the first trial in adaptation (dark red, Ad) and readaptation blocks (dotted red, Re) across all subjects. The mean baseline EMG activity (reach trials without force field in baseline block) is in magenta. Traces are aligned on reach onset (t=0, vertical brown line). The dotted grey rectangular box represents 100 ms window before reach onset. b) Normalized EMG activity in posterior deltoid muscle during CCW perturbation (group 1) in adaptation (dark blue, Ad) and readaptation blocks (dotted blue, Re) averaged across all subjects. The mean baseline EMG activity (reach trials without force field in baseline block) is in green. Note that a and b are representative graphs to show the temporal evolution of EMG and similar graphs for G2 CW DELT and G1 CCW PEC are not shown for simplicity. c) Mean EMG activity during the 100ms window (−100 to 0) before reach onset for muscles PEC and DELT during baseline (Ba), adaptation (Ad) and readaptation (Re) blocks in both groups G1 and G2. Each dot represents the first trial in the corresponding block. The black asterisks (*) denote the mean across all participants (i.e., both CW and CCW groups). For both muscles, the mean EMG activity before reach onset is similar in all the blocks, which nullifies any possibility of coactivation strategy during the reaching movements of the readaptation block.

To summarize the Results, we first reproduced the expression of savings as faster relearning in human reach adaptation. We found evidence that the learning curves across trials converged faster to their asymptote in the readaptation blocks based on the pathlengths and the correlations between applied and measured forces. Building on these results our original contribution was to demonstrate that the very first trial in the readaptation block possessed all the characterizing features of better-adapted movements. Since the end of the washout was similar to the baseline performance, as highlighted by similar initial angles, pathlengths, positional variances, and the fact that participants were not aware that there would be a second adaptation phase, the contribution of an explicit strategy or feedforward adaptation could both be ruled out. Moreover, the readaptation phase started with similar levels of background muscle activity, and we even observed similar initial angles suggesting comparable effects of the unexpected force field. Yet, the kinematic and kinetics markers of better-adapted trials were present in the very first relearning trials: smaller path length, lateral deviation, target overshoot, peak terminal force and the higher correlation between commanded and applied forces. Thus, the improvement observed in the very first relearning trial can be explained by feedback adaptation, and the associated improvements contributed to the faster convergence towards the asymptote of the learning curve across trials.

## Discussion

We documented the presence of a feedback adaptation component in force field reaching movements during readaptation to a force field previously encountered. Healthy participants first adapted to a unidirectional force field, followed by a washout sufficiently long to ensure that trials recovered a straight path, and then re-exposure to the same force field to quantify the nature of readaptation. First, we confirmed that readaptation was faster than adaptation based on the observation that the time to reach the asymptote of the learning curve in the readaptation block was significantly shorter than that in the adaptation block. This observation was consistent with previous findings (Smith et al., 2006; Herzfeld et al., 2014; Stockinger et al., 2014; Coltman et al., 2019) and confirmed the presence of savings (Shadmehr and Brashers-Krug, 1997; Overduin et al., 2006).

Second, in addition to the faster relearning rate characterizing trial-by-trial changes in motor performances, our main contribution was to highlight that, not only were participants relearning faster, but the very first trial of the readaptation phase was already better adapted. We observed a reduced pathlength, target overshoot, lateral deviation, terminal peak force and a higher commanded-measured force correlation in comparison with the first trials of the adaptation phase, despite the absence of anticipation in the relearning phase, and without any measurable change in average muscle activity that could have modulated limb stiffness (Burdet et al., 2001). This supports the hypothesis that an efficient feedback control response was recalled during the very first readaptation trial and improved stabilization of the ongoing movement. We believe that the link between feedback adaptation and savings is direct, considering that the state of adaptation at the end of the first relearning trial carries over to the next, which in turn enables better performance and feedback corrections for the following trials. These observations demonstrate that the neural mechanism underlying faster relearning involves a feedback adaptation component. They also provide further evidence that feedback adaptation plays a central role in the traditionally called feedforward component associated with trial-by-trial sequential adaptation scenarios (Crevecoeur et al., 2020b; Mathew et al., 2020).

In the context of adaptation to visual disturbances such as a cursor rotation, the presence of savings has been debated due to the presence of both implicit and explicit strategies (Taylor et al., 2014; McDougle et al., 2015; Morehead et al., 2015), the latter consisting in re-aiming at a remapped target. It is important to note that the possibility to handle disturbances with an explicit re-aiming strategy is potentially specific to visuomotor rotations (although subject to interference (Mazzoni and Krakauer, 2006)). It is less clear that such an explicit strategy can produce apparent savings during adaptation to a force field, which involves a time-varying force that cannot be fully compensated simply by changing the aiming direction. In our experiment, washout was complete, participants did not anticipate the relearning session and we concentrated on the very first relearning trial to rule out any influence of such an explicit strategy. Note that none of the previous studies explicitly investigated the first relearning trial in detail with multiple kinematic variables (Fig 3) to highlight the improvement in relearning (Huberdeau et al., 2015; Leow et al., 2016). As a consequence and based on metrics specific to anticipation (for instance the initial angle), one may miss the properties of the first relearning trial that become apparent when considering the whole trace, including later within-trial corrections. We suggest that a feedback adaptation component recalling a previously acquired representation during movement produced a clear improvement during the very first relearning trial. Interestingly, previous studies documented similar adaptive control strategies when exposed to random visuomotor rotations (Braun et al., 2009), thus there remain open questions about the possibility that feedback adaptation also influences savings in visuomotor learning.

Increasing evidence provides a compelling indication that the feedback control system plays a central role in motor adaptation. Recent studies highlighted features of the feedback adaptation component such that the feedback response was specifically and finely tuned to the ongoing force field perturbation, not only across trials with the same force field but also across different kinds of force fields (Crevecoeur et al., 2020a). The underlying mechanism associated with feedback adaptation can be preserved in memory long enough to contribute to the trial-by-trial adaptation scenarios (Mathew et al., 2020). Also, the changes in muscle activity consistent with feedback adaptation occurred within about 250ms following reach onset, which is proposed as an estimate of the latency of motor adaptation in the nervous system (Crevecoeur et al., 2020b, 2020a). It was previously highlighted that modulation of long-latency feedback parallels the time course of the fast learning timescale (Coltman et al., 2019; Coltman and Gribble, 2020), which very likely is a major contributor to the speeding up of adaptation given the short time scale at which the faster relearning occurs. We add to this line of evidence by showing that feedback adaptation underlies savings in human force field learning, as it is clear that the behavioural improvement in the first relearning trial is attributable to the feedback control system, with the possibility that all feedback mechanisms are engaged (short- or long-latency reflexes, and visually-mediated corrections) (Franklin and Wolpert, 2008; Cross et al., 2019).

In conclusion, we suggest that there is a genuine acceleration of learning in a readaptation context that is supported by adaptive feedback control. These results imply one challenging question for future work, that is to disentangle the contribution of feedback adaptation and explicit re-aiming strategies in the overall speedup of trial-by-trial learning across paradigms.

## Data Availability

Study data and codes used in this project are available at the following link: https://zenodo.org/record/4570064#.YDzv67hKhPY

## Acknowledgements

This work was funded by FRS-FNRS PDR grant T.0048.19 (Belgium).

